# Autophagy Compensates for Lkb1 Loss to Maintain Adult Mice Homeostasis and Survival

**DOI:** 10.1101/2020.08.26.268003

**Authors:** Khoosheh Khayati, Vrushank Bhatt, Zhixian Sherrie Hu, Sajid Fahumy, Xuefei Luo, Jessie Yanxiang Guo

## Abstract

Liver Kinase B1 (LKB1) is the major energy sensor for cells to respond to metabolic stress. Autophagy degrades and recycles proteins, macromolecules, and organelles for cells to survive starvation. To access the role and cross-talk between autophagy and Lkb1 in normal tissue homeostasis, we generated genetically engineered mouse models where we can conditionally delete *Lkb1* and autophagy essential gene, *Atg7, respectively or simultaneously*, throughout the adult mice. We found that Lkb1 was essential for the survival of adult mice, and autophagy activation could temporarily compensate for the acute loss of Lkb1 and extend mouse life span. We further found that acute deletion of Lkb1 in adult mice led to impaired intestinal barrier function, hypoglycemia, and abnormal serum metabolism, which was partly rescued by the Lkb1 loss-induced autophagy upregulation via inhibiting p53 induction. Taken together, we demonstrated that autophagy and Lkb1 work synergistically to maintain adult mouse homeostasis and survival.

## Introduction

Liver kinase B1 (LKB1) is a tumor suppressor, metabolic sensor and master modulator of AMP-activated protein kinase (AMPK) and mammalian target of rapamycin complex1 (mTORC1) activity, leading to the control of energy metabolism, cell polarity, cell survival and proliferation (1-3). Heterozygous germline mutations in *LKB1* lead to the development of Peutz-Jeghers syndrome (PJS), an autosomal dominant disease with hamartomatous polyp formation in the gastrointestinal tract (4). Constitutive deficiency of Lkb1 leads to embryonic lethality due to impaired neural tube closure and somitogenesis, mesenchymal tissue cell death, and defective vasculature (5). Specific deletion of Lkb1 in vascular endothelial cells results in dilated embryonic vessels and death at E12.5, which is attributed to the reduced Tgfβ signaling in yolk sac (1, 6). Liver-specific deficiency of Lkb1 causes impaired glucose metabolism (7). Muscle-specific deletion of Lkb1 results in lower fasting blood glucose and insulin levels, along with increased glucose uptake through muscles (8). Lkb1 loss in hematopoietic stem cells causes dysfunctional mitochondria, leading to pancytopenia due to reduced levels of ATP, fatty acids and nucleotides (3, 9, 10). Taken together, tissue-specific knockout studies underscore the importance of Lkb1 in tissue homeostasis, metabolism, and stem cell maintenance. Somatic *LKB1* mutations are related with a number of human cancers; however, tissue-specific removal of *Lkb1* in mice does not necessarily lead to tumor formation (11).

Autophagy, a highly conserved self-degradative process, plays an essential role in cellular stress responses and survival (12-14). Yeast cells rely on autophagy to survive nitrogen starvation (15); neonatal mice depend on autophagy to survive neonatal starvation-induced amino acid depletion (12); and adult mice requires autophagy to survive starvation (14, 16).

Given that both Lkb1 signaling and autophagy play indispensable roles in maintaining tissue energy homeostasis, we began to investigate the interaction of Lkb1 signaling and autophagy in supporting homeostasis of adult mice. We engineered mice to conditionally (Tamoxifen (TAM)-inducible) and systemically delete *Lkb1* and *Atg7*, either respectively or simultaneously. Same as previous report (16), systemic *Atg7* ablation led to extensive liver and muscle damage, and neurodegeneration starting at 6 weeks post-deletion, and limited mouse survival to 3 months. Surprisingly, we found that adult mice with acute ablation of *Lkb1* through whole-body (*Lkb1*^*-/-*^ mice) died within 30 days and showed upregulated autophagy in most tissues. Moreover, systemic co-deletion of *Lkb1* and *Atg7* limited mice survival to 20 days. *Lkb1*^*-/-*^ mice displayed disruption of intestinal structure and impaired intestinal defense barrier, which was deteriorated by co-deletion with *Atg7*. Supplementation of broad-spectrum antibiotics or systemic deletion of *p53* partly rescued the death of the mice with concurrent deletions of *Lkb1* and *Atg7*, but not the mice with *Lkb1* deletion alone. Serum metabolomics profiling analysis showed that acute short-term deletion of *Atg7* or *Lkb1* respectively significantly decreased the levels of most essential and non-essential amino acids and some metabolites involved in the tricarboxylic acid (TCA) cycle, urea cycle and glycolysis. This phenotype was further enhanced in mice with concurrent deletions of *Atg7* and *Lkb1*. Taken together, this study reveals a novel role of Lkb1 in the management of tissue homeostasis in adult mouse and the mechanism by which autophagy upregulation temporarily compensates for the acute loss of Lkb1.

## Results

### Acute systemic *Lkb1* deletion upregulates autophagy in adult mouse

Tissue-specific *Lkb1* knockout studies demonstrate that Lkb1 plays an important role in supporting tissue and organ homeostasis (11). However, how Lkb1 regulates adult mouse homeostasis is unknown. Autophagy is required to maintain tissue homeostasis and mouse survival in starvation (14, 16). Whether and how Lkb1 and Atg7 interact to maintain tissue homeostasis for adult mouse survival remains an open question. To address this, we generated genetically engineered mouse models, in which *Atg7* and *Lkb1* were surrounded by flox alleles, whereas the expression of a TAM-regulated Cre-recombinase was manipulated through the ubiquitously expressed ubiquitin C (Ubc) promoter (17). Four mouse strains were generated: *Ubc-CreERT2*^*/+*^, *Ubc-CreERT2*^*/+*^;*Atg7*^*flox/flox*^, *Ubc-CreERT2*^*/+*^;*Lkb1*^*flox/flox*^, *and Ubc-CreERT2*^*/+*^;*Atg7*^*flox/flox*^;*Lkb1*^*flox/flox*^. Following TAM injections in 8-10 week-old mice, Cre is activated, leading to the deletion of *Atg7* or *Lkb1*, and producing a near complete and sustained loss of Atg7 protein (*Atg7*^*-/-*^ mice), Lkb1 protein (*Lkb1*^*-/-*^ mice), or dual deletions of Atg7 and Lkb1 proteins (*Atg7*^*-/-*^;*Lkb1*^*-/-*^ mice) in all tissues (Fig. 1a). The deletions of Lkb1 and Atg7 throughout the mouse tissues were confirmed by western blot (Fig. 1b). In addition, accumulation of autophagy substrate p62 was observed in all tissues of the mice with Atg7 ablation (*Atg7*^*-/-*^ mice and *Atg7*^*-/-*^;*Lkb1*^*-/-*^ mice) (Fig. 1c), indicating autophagy blockade.

**Fig 1.**
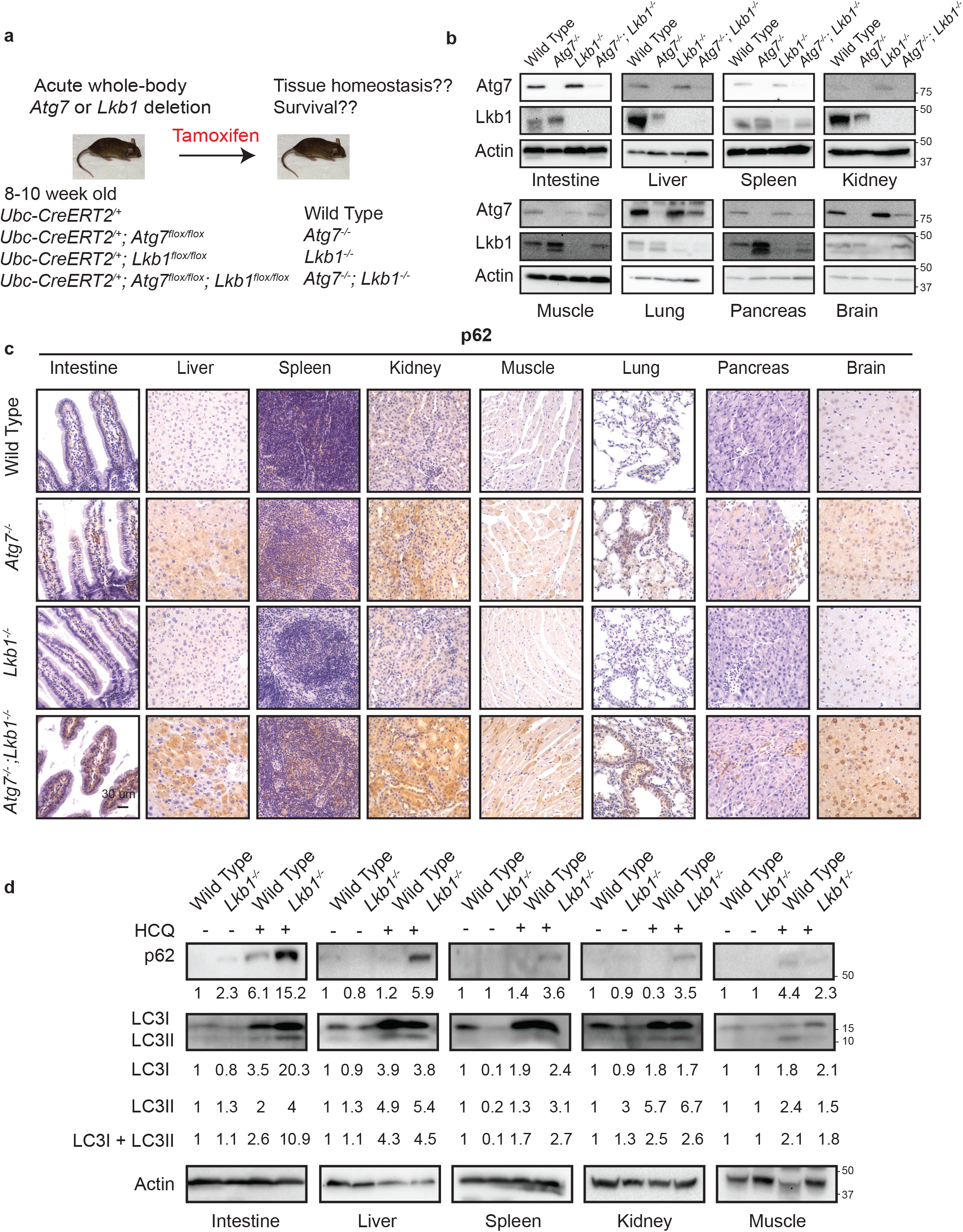
Autophagy is up-regulated in tissues of *Lkb1-*deficient mice. a. Experimental design for generation of *Atg7*^*-/-*^, *Lkb1*^*-/-*^, *and Atg7*^*-/-*^;*Lkb1*^*-/-*^ mice. b. Western blotting for Atg7 and Lkb1 of the indicated tissues from WT control, *Atg7*^*-/-*^, *Lkb1*^*-/-*^, *and Atg7*^*-/-*^;*Lkb1*^*-/-*^ adult mice. β-actin serves as a protein loading control. c. Representative IHC for p62 of different tissues from WT control, *Atg7*^*-/-*^, *Lkb1*^*-/-*^, *and Atg7*^*-/-*^;*Lkb1*^*-/-*^ adult mice. d. Western blotting for p62, LC3I and LC3II from different tissues of WT control and *Lkb1*^*-/-*^ mice with or without HCQ treatment represents the up-regulation of autophagy in *Lkb1-* deficient mice. β-actin serves as a protein loading control. Numbers indicate the quantification of protein levels normalized to actin and WT control.

Given that AMPK is required for autophagy activation, Lkb1 deficiency leads to the loss of AMPK activity and suppression of autophagy (18). Consequently, studying the role of autophagy in the mice with systemic loss of Lkb1 may be considered counterintuitive. However, studies from us and other groups have shown that AMPK is activated in the mouse *Kras*-driven lung tumors with Lkb1 loss (19, 20). Indeed, we also observed that autophagy is required for *Kras*-mutant *Lkb1*-deficient lung tumorigenesis (19). Instead of Lkb1, AMPK can be activated by calmodulin-dependent kinase kinase (CaMKK), and transforming growth factor beta-activated kinase1 (TAK1) (21, 22), further leading to autophagy activation. We therefore examined the autophagy status in *Lkb1*^*-/-*^ mice by administering the mice with hydroxychloroquine (HCQ) to block autophagy flux. Accumulation of autophagy substrates including both microtubule-associated protein 1A/1B-light chain3 (LC3) I and II along with p62 were observed in most tissues of mice after 5 hours of HCQ treatment. Moreover, HCQ treatment led to a higher accumulation of LC3-II and p62 in *Lkb1*-deficient mice compared with wild type (WT) control mice (Fig. 1d), indicating that autophagy is activated and upregulated in the mice with acute loss of Lkb1.

### Interaction of Lkb1 and autophagy is required for adult mouse survival

It has previously been shown that mice with systemic autophagy ablation have a life span of 2-3 months and the mortality is due to initial Streptococcus infection and eventual neurodegeneration (16). Whereas, adult mice with whole-body *Lkb1* deletion survive for up to 6 weeks (23). Given that both Lkb1 signaling and autophagy pathway regulate cellular homeostasis (12, 16, 18, 24), we hypothesize that upregulation of autophagy in *Lkb1-*deficient mice might compensate for the acute Lkb1 loss to maintain energy homeostasis for mouse survival. Therefore, we evaluated the overall survival of adult mice with acute deletion of *Atg7* and *Lkb1* respectively or in combination. Our data reproduced the survival rate for *Atg7-*deficient mice when *Atg7* was acutely deleted in adult mice (Fig. 2a) (16). The life span of *Lkb1-*deficient mice was limited to 4 weeks post-TAM administration. Co-deletion of *Atg7* and *Lkb1* led to a significant decrease in the survival of *Atg7*^*-/-*^;*Lkb1*^*-/-*^ mice compared with *Lkb1*^*-/-*^ or *Atg7*^*-/-*^ mice, resulting in less than 3 weeks of survival (Fig. 2a).

**Fig 2.**
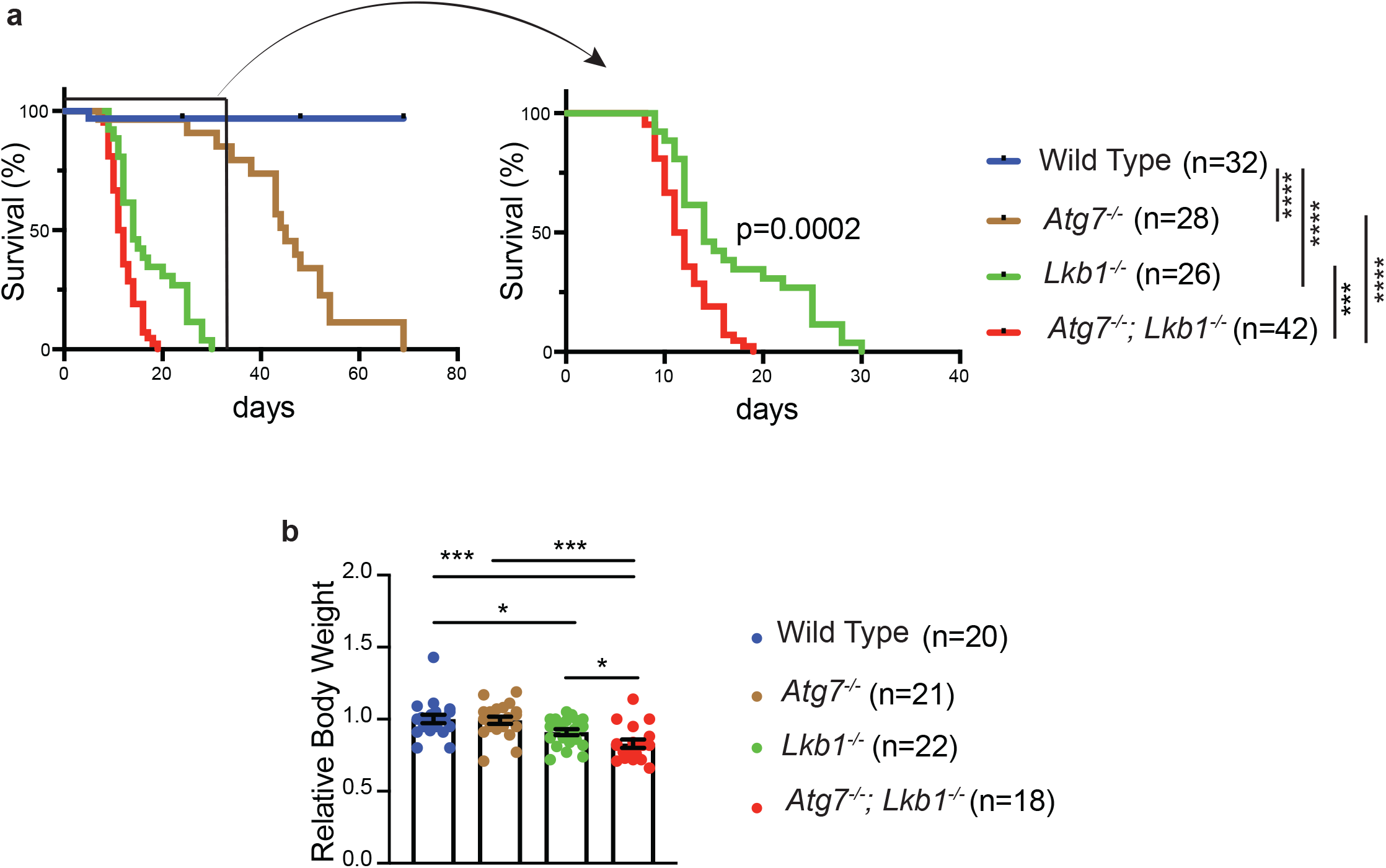
Autophagy compensates for acute Lkb1 loss to support the survival of adult mice. a. Kaplan-Meier survival curve of WT control, *Atg7*^*-/-*^, *Lkb1*^*-/-*^, *and Atg7*^*-/-*^;*Lkb1*^*-/-*^ adult mice. *** P<0.001, and ****P<0.0001 (log-rank Mantel-Cox test). b. Relative body weights of WT control, *Atg7*^*-/-*^, *Lkb1*^*-/-*^, *and Atg7*^*-/-*^;*Lkb1*^*-/-*^ adult mice normalized to the body weight of WT control mice. Data are mean ± s.e.m. *P<0.05, *** P<0.001.

At 15 days post-TAM administration, compared with WT control mice, there was a significant loss of body weight in *Lkb1*^*-/-*^ mice, which was further deteriorated by the loss of Atg7 (Fig. 2b). Hematopoietic-specific *Lkb1*-deficient mice died from pancytopenia (9). We did not observe any differences of red and white blood cell and platelet count among WT control, *Atg7*^*-/-*^, *and Lkb1*^*-/-*^ mice; however, the platelet count in *Atg7*^*-/-*^;*Lkb1*^*-/-*^ mice was significantly higher than that in WT control mice (Supplementary Fig. 1).

### Acute autophagy ablation aggravates Lkb1-deficiency-induced loss of secretory cell structure in small intestine

To elucidate the underlying mechanism by which *Atg7* and *Lkb1* ablation alone or in concurrent impacts the mouse survival, we examined the histology of different tissues. After short-period (15 days) of protein deletion, which is before the death of *Atg7*^*-/-*^;*Lkb1*^*-/-*^ mice, the damage of tissues was not observed in *Atg7*-deficient mice (Supplementary Fig. 2a). Moreover, except for the intestine (Fig. 3a), most of the tissues in both *Lkb1*^*-/-*^ and *Atg7*^*-/-*^;*Lkb1*^*-/-*^ mice were not visibly affected as examined by hematoxylin and eosin (H&E) staining (Supplementary Fig. 2a). Same as tissue-specific *Lkb1* deletion in intestinal-epithelium cells (25), enlarged undifferentiated goblet-Paneth cells in the crypt of intestine, including duodenum, jejunum, and ileum, were observed in *Lkb1*^*-/-*^ mice, which was further exacerbated by the concurrent ablation of *Atg7* in *Atg7*^*-/-*^;*Lkb1*^*-/-*^ mice (Fig. 3a). This observation was further confirmed by Alcian blue staining for goblet cells and immunohistochemistry (IHC) for lysozyme staining to detect Paneth cells (Fig. 3b and c). The decreased intensity of lysozyme staining indicates less frequent and undifferentiated Paneth cells in *Atg7*^*-/-*^;*Lkb1*^*-/-*^ mice (Fig. 3c). In addition to western blot (Fig. 1b), the deletion of Lkb1 and Atg7 specifically in intestine was also confirmed by IHC (Supplementary Fig. 2b). In consistent with induction of autophagy in *Lkb1*-deficient mice, LC3-II puncta were observed in the intestine of *Lkb1*^*-/-*^ mice (Supplementary Fig. 2c).

**Fig 3.**
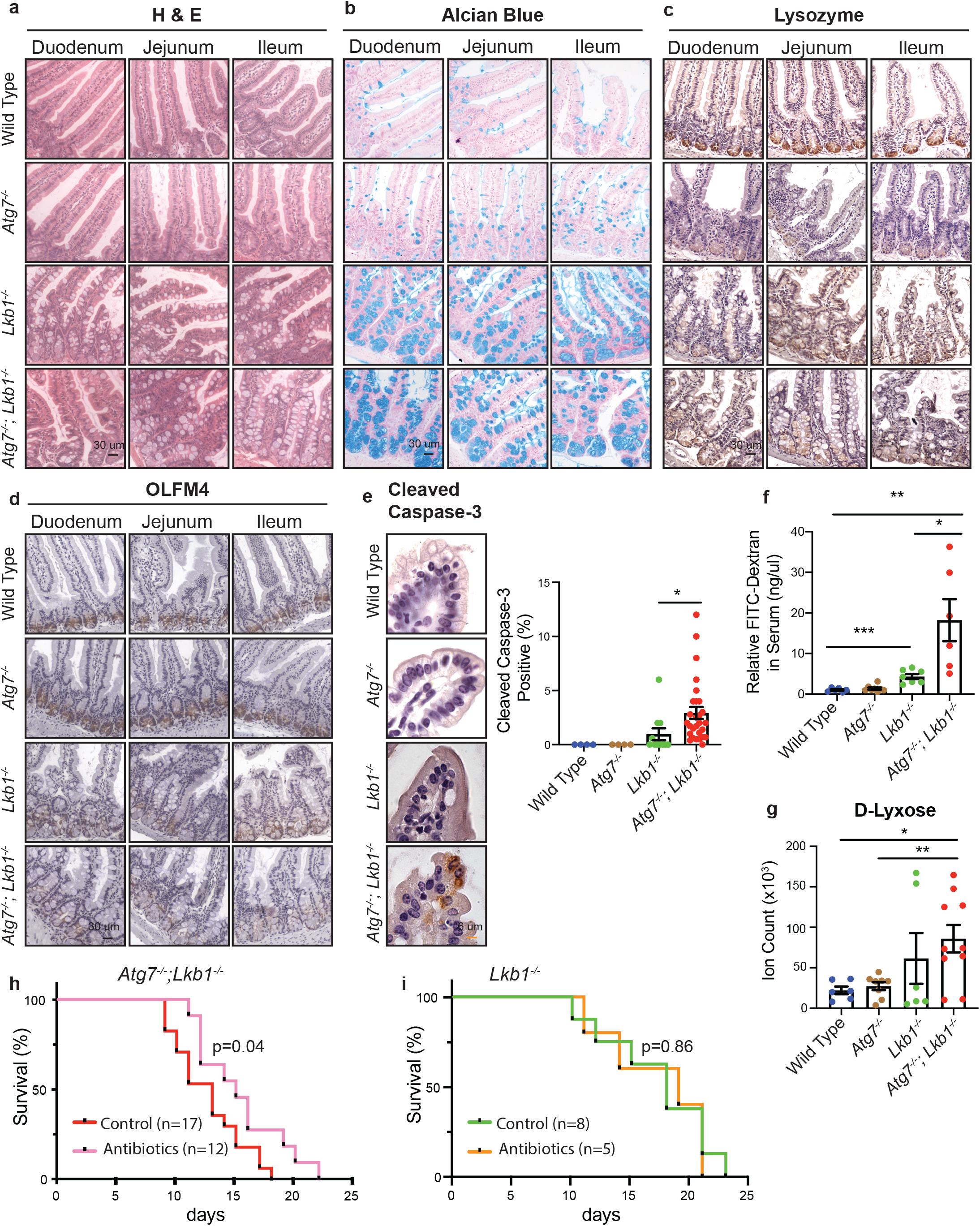
Autophagy ablation deteriorates impaired intestinal structure and function caused by acute *Lkb1* deletion. a. Representative H&E staining of duodenum, jejunum, and ileum for WT control, *Atg7*^*-/-*^, *Lkb1*^*-/-*^ and *Atg7*^*-/-*^;*Lkb1*^*-/-*^ adult mice. b. Alcian blue staining of adult mouse intestine shows the enlargement of mucin-secreting cells in *Lkb1*^*-/-*^ and *Atg7*^*-/-*^;*Lkb1*^*-/-*^ mice. c. Representative IHC for intestinal lysozyme shows decrease of Paneth cell population in *Lkb1*^*-/-*^ and *Atg7*^*-/-*^;*Lkb1*^*-/-*^ crypts. d. Representative IHC for OLFM4 of intestine shows the decrease of stem cells with greater extent in *Atg7*^*-/-*^;*Lkb1*^*-/-*^ crypts compared with WT control and *Lkb1*^*-/-*^ mice. e. Left: Representative IHC for cleaved caspase-3 of intestine delineates increase of cell death in *Atg7*^*-/-*^;*Lkb1*^*-/-*^ at tips of villi compared with WT control and *Lkb1*^*-/-*^ mice. Right: Quantification of cleaved caspase-3. Data are mean ± s.e.m. *P<0.05. f. Representative relative FITC-dextran levels in sera of WT control, *Atg7*^*-/-*^, *Lkb1*^*-/-*^ and *Atg7*^*-/-*^;*Lkb1*^*-/-*^ adult mice at 4 hours post-oral gavage of FITC-dextran. Data are mean ± s.e.m. *P<0.05, **P<0.01, ***P<0.001. g. The level of serum D-lyxose in WT control, *Atg7*^*-/-*^, *Lkb1*^*-/-*^ and *Atg7*^*-/-*^;*Lkb1*^*-/-*^ adult mice measured by LC-MS shows an increase of D-lyxose in *Atg7*^*-/-*^;*Lkb1*^*-/-*^ sera compared with WT control mice. Data are mean ± s.e.m. *P<0.05, **P<0.01. h. Kaplan-Meier survival curve of *Atg7*^*-/-*^;*Lkb1*^*-/-*^ mice treated without or with broad-spectrum antibiotics (log-rank Mantel-Cox test). i. Kaplan-Meier survival curve of *Lkb1*^*-/-*^ mice treated without or with broad-spectrum antibiotics (log-rank Mantel-Cox test).

To evaluate the role of Lkb1 and autophagy in other components of the small intestinal crypt, we examined the status of stem cells that reside at the bottom of the crypt for regenerating almost all the epithelium cells, including Paneth and goblet cells, enterocytes and tuft cells (26, 27). IHC for olfactomedin4 (OLFM4) (an intestine stem cell marker) shows the lower intensity and frequency of stem cells in *Lkb1*^*-/-*^ mice compared with WT control or *Atg7*^*-/-*^ mice (Fig. 3d). Moreover, co-deletions of *Lkb1* and *Atg7* further reduced the intensity and frequency of the cells expressing OLFM4 compared with *Lkb1* deletion alone (Fig. 3d). However, short-term deletion of *Atg7* alone only impaired Paneth cell formation (Fig. 3c) (28).

The structure of the small intestine was extremely damaged by *Lkb1* deletion alone or co-deletions of *Lkb1* and *Atg7* (Fig. 3a-d), which could be due to less regeneration from intestinal stem cells or increased cell death. Cell death in small intestine occurs through apoptosis at the tip of the villi which eventually leads to shedding of dead cells into the lumen (29, 30). We found that there was a significant increase of apoptotic cell death in the tip of the villi in *Atg7*^*-/-*^;*Lkb1*^*-/-*^ mice compared with *Lkb1*^*-/-*^ mice (Fig. 3e). Intestinal crypt is the region for cell division and migration to upper sites of the villi (31). Compared with the WT control mice, we did not observe any significant difference in the cell proliferation rate in the crypt of the mice lacking either *Atg7* or *Lkb1* alone or in combination (Supplementary Fig. 2d).

Taken together, we show that Lkb1 is necessary for maintaining the structural integrity of the intestine and that autophagy activation partly compensates for the severe intestinal phenotype induced by the loss of Lkb1.

### Autophagy activation in *Lkb1*-deficient mice protects the intestinal epithelium-barrier function

Intestinal epithelium cell-specific deletion of *Lkb1* results in an increased susceptibility to dextran sodium sulfate-induced colitis and a definitive shift in the composition of the microbial population in the mouse intestine (32), suggesting that Lkb1 plays an important role in maintaining the immune barrier function of the intestinal epithelium. Moreover, it has recently been reported that autophagy is essential for the maintenance of Lgr5^+^ stem cells and regeneration of epithelium barrier during cytotoxic stress (33). We therefore examined the integrity of intestinal epithelium-barrier in mice with systemic deletion of *Atg7* or *Lkb1* alone, or their co-deletion by measuring the Fluorescein Isothiocyanate (FITC)-dextran in-fluxed from the gastrointestinal tract to peripheral circulation. We observed significantly increased levels of serum FITC-dextran in *Lkb1*-deficient mice compared with WT control mice, which is further increased by the co-deletion with *Atg7* in *Atg7*^*-/-*^;*Lkb1*^*-/-*^ mice (Fig. 3f). However, short-term systemic ablation of *Atg7* alone did not impair intestinal epithelium-barrier (Fig. 3f). This observation suggests that upregulated autophagy by acute *Lkb1* deletion is required for the maintenance of intestinal epithelium-barrier.

Loss of intestinal epithelium-barrier causes mice to be susceptible to bacterial infection (32). We found that the levels of D-lyxose, an aldopentose sugar and a component of the bacterial glycolipids (34), was significantly increased in *Atg7*^*-/-*^;*Lkb1*^*-/-*^ mice compared with WT control (Fig. 3g), indicating defective bacterial defense. We therefore hypothesized that increased bacterial infection by loss of intestinal epithelium barrier could contribute to the mouse death caused by co-deletions of *Lkb1* and *Atg7*. Hence, we treated the *Lkb1*^*-/-*^ and *Atg7*^*-/-*^;*Lkb1*^*-/-*^ mice with broad-spectrum antibiotics and assessed the mouse survival rate in comparison with untreated ones. Broad-spectrum antibiotics administration significantly extended the survival of *Atg7*^*-/-*^;*Lkb1*^*-/-*^ mice (Fig. 3h), whereas, did not affect the lifespan of *Lkb1*^*-/-*^ mice (Fig. 3i). Thus, one of the potential mechanisms of autophagy activation in response to acute systemic Lkb1 deficiency could be to maintain the survival of mice by preventing bacterial invasion.

### Autophagy activation in *Lkb1*-deficient mice prevents p53 activation to maintain mouse survival

Given that autophagy drives an inhibitory role towards p53 activation (12, 35, 36), we expected to observe an increased p53 activation in autophagy-deficient mouse tissues. Indeed, IHC staining for p53 showed that the frequency of nuclear p53 was significantly higher in most tissues of *Atg7*^*-/-*^;*Lkb1*^*-/-*^ mice compared with *Lkb1*^*-/-*^ mice or WT control mice after short-term deletion of the genes (Fig. 4a and Supplementary Fig. 3). This is also accompanied by the significantly increased mRNA levels of p53-targted downstream genes such as *p21* and phosphatase and tensin homolog (*PTEN*) in *Atg7*^*-/-*^;*Lkb1*^*-/-*^ mice compared with *Lkb1*^*-/-*^ mice or WT control mice (Fig. 4b). Accordingly, we tested the hypothesis that activation of autophagy by acute *Lkb1* ablation could prevent mouse death by inhibiting p53 activation. To address this, two new cohorts of mice were generated: *Ubc-CreERT2*^*/+*^;*Lkb1*^*flox/flox*^;*p53*^*flox/flox*^, *and Ubc-CreERT2*^*/+*^;*Atg7*^*flox/flox*^;*Lkb1*^*flox/flox*^;*p53*^*flox/flox*^. TAM administration can cause concurrent deletion of *Lkb1* and *p53* (*Lkb1*^*-/-*^;*p53*^*-/-*^ mice) or *Atg7, Lkb1* and *p53* (*Atg7*^*-/-*^;*Lkb1*^*-/-*^;*p53*^*-/-*^ mice) throughout the whole body. We evaluated the role of *p53* deletion in mouse survival by comparing with *Lkb1*^*-/-*^ mice and *Atg7*^*-/-*^;*Lkb1*^*-/-*^ mice that have intact *p53*. Systemic *p53* ablation had no effect on the survival of *Lkb1*-deficient mice. However, whole-body ablation of *p53* significantly extended the life span of *Atg7*^*-/-*^;*Lkb1*^*-/-*^ mice (Fig. 4c). Thus, when *Lkb1* is acutely ablated throughout the adult mice, autophagy inhibits p53 activation to temporarily extend the mouse life span.

**Fig 4.**
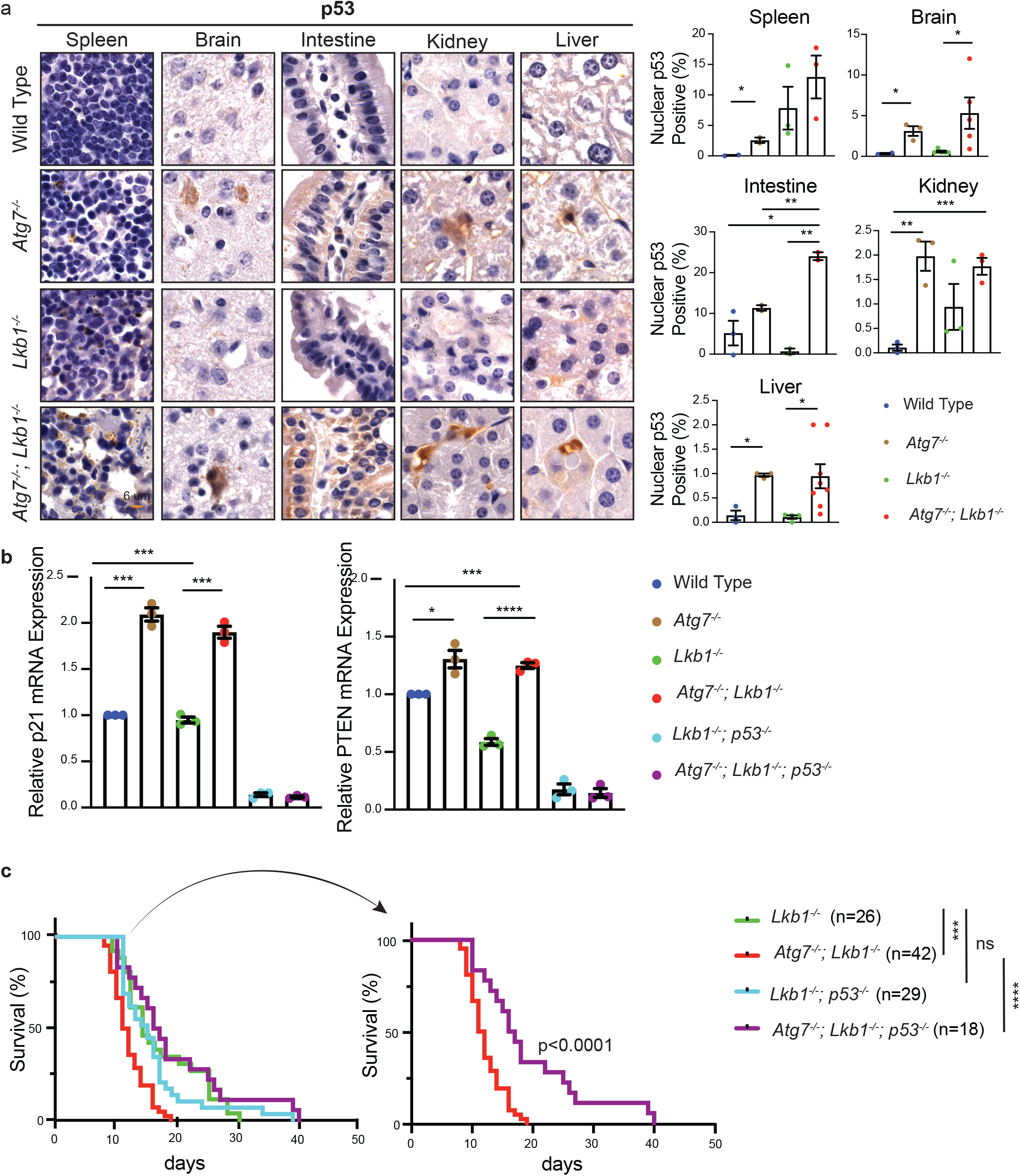
p53 deficiency extends the life span of *Atg7*^*-/-*^;*Lkb1*^*-/-*^ mice. a. Left: Representative IHC for p53 in different tissues of WT control, *Atg7*^*-/-*^, *Lkb1*^*-/-*^ and *Atg7*^*-/-*^;*Lkb1*^*-/-*^ adult mice shows an increase of nuclear p53 in *Atg7*-ablated tissues. Right: Bar graphs represents the quantification of nuclear p53. Data are mean ± s.e.m. *P<0.05, **P<0.01, and ***P<0.001. b. Quantitative real-time PCR of *Cdkn1a* (*p21*) and *PTEN* for intestine tissues of WT control, *Atg7*^*-/-*^, *Lkb1*^*-/-*^ and *Atg7*^*-/-*^;*Lkb1*^*-/-*^ adult mice. Data are mean ± s.e.m. *P<0.05, ***P<0.001, and ****P<0.0001. c. Kaplan-Meier survival curve of *Lkb1*^*-/-*^, *Atg7*^*-/-*^;*Lkb1*^*-/-*^, *Lkb1*^*-/-*^;*p53*^*-/-*^ and *Atg7*^*-/-*^;*Lkb1*^*-/-*^;*p53*^*-/-*^ adult mice. ***P<0.001, ****P<0.0001 and ns: non-significant (log-rank Mantel-Cox test).

### Lkb1 and Atg7 are required to maintain adult mice homeostasis

To further elucidate the underlying mechanism of Lkb1 and Atg7 in supporting adult mice survival, we performed serum metabolomics in fasting state after short-term deletion of the genes. Of the 90 metabolites we examined, we found that acute deletion of *Atg7* or *Lkb1* alone significantly decreased the levels of most essential and non-essential amino acids and some metabolites involved in urea cycle and glycolysis (Fig. 5a-d). Interestingly, we found that the reduced levels of TCA cycle intermediates were only observed in the absence of *Atg7* (*Atg7*^*-/-*^ mice and *Atg7*^*-/-*^; *Lkb1*^*-/-*^ mice), but not in *Lkb1*-deficient mice (Fig. 5b). We further found that in both fasted state (Fig. 5d) and fed state (Fig. 5e), blood glucose level was significantly lower in *Lkb1*^*-/-*^ and *Atg7*^*-/-*^; *Lkb1*^*-/-*^ mice compared with WT control mice. Following that, we evaluated the serum insulin levels in all 4 groups of mice. The insulin levels in *Lkb1*^*-/-*^ and *Atg7*^*-/-*^;*Lkb1*^*-/-*^ mice were decreased with the same trend as glucose compared with WT control mice (Fig. 5f), delineating the possibility of pancreatic malfunction in these mice, although the damage in pancreas was not visible by histology (Supplementary Fig. 2a). We also performed a metabolomics profiling analysis of the intestinal tissue (ileum) that showed significantly impaired histology and function in both *Lkb1*^*-/-*^ and *Atg7*^*-/-*^;*Lkb1*^*-/-*^ mice (Fig. 3). The alteration of metabolic pathways in the intestine metabolomics profiling due to *Atg7* and *Lkb1* ablation was consistent with the change in the serum metabolomics profiling, ie, the loss of *Lkb1* alone or together with *Atg7* ablation resulted in the decreased levels of certain intermediates involved in the amino acid metabolism, TCA cycle, urea cycle and glycolysis (Supplementary Fig. 4). Thus, autophagy synergizes with Lkb1 to maintain host homeostasis in the adult mice.

**Fig 5.**
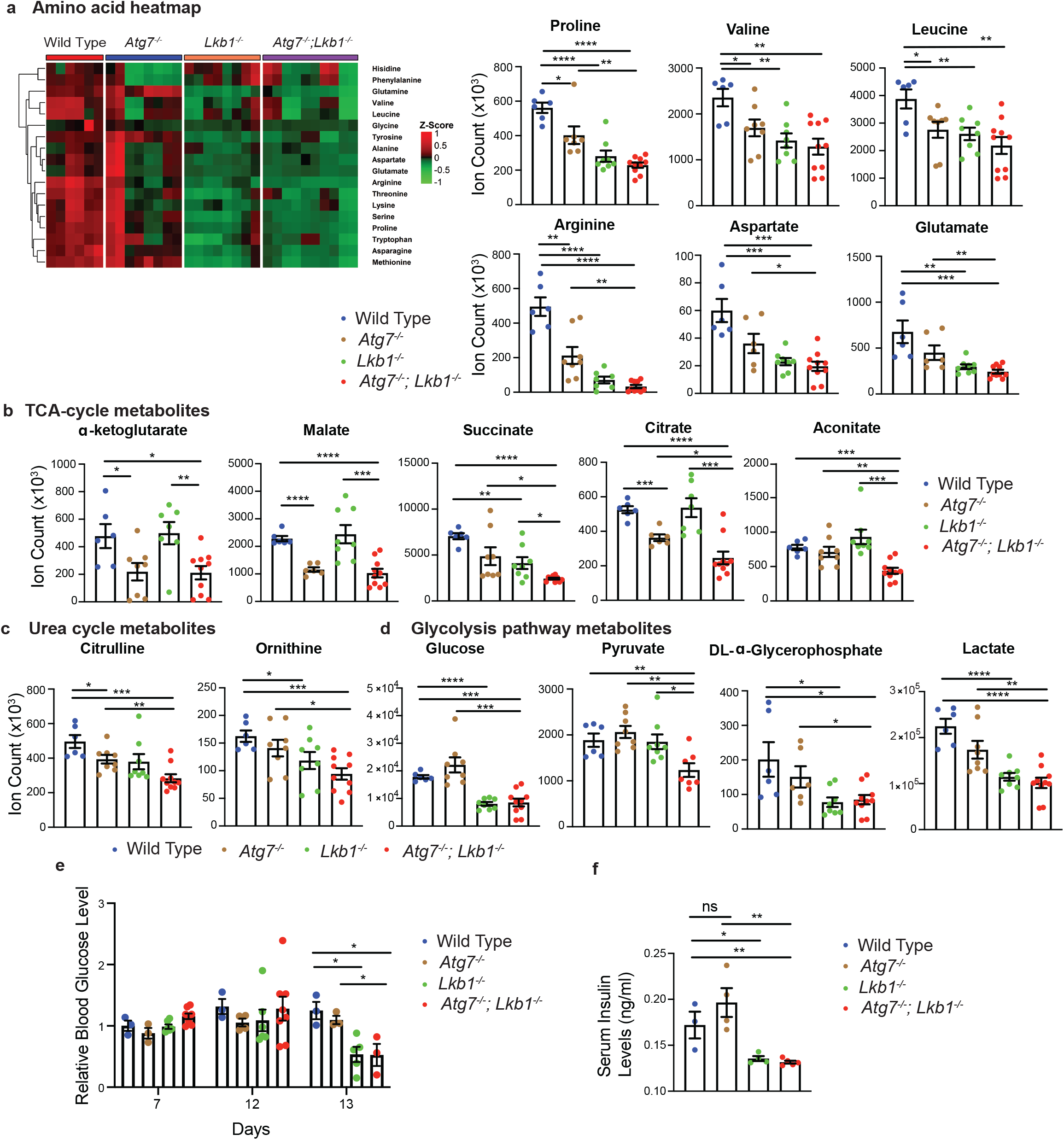
Major metabolic pathways are disturbed in *Lkb1*^*-/-*^ and *Atg7*^*-/-*^;*Lkb1*^*-/-*^ mice. a. Left: Representative heat map of all the amino acids in sera of WT control, *Atg7*^*-/-*^, *Lkb1*^*-/-*^ and *Atg7*^*-/-*^;*Lkb1*^*-/-*^ adult mice compared with WT control mice at fasting state. Right: Bar graphs show the levels of amino acids that are significantly decreased in *Lkb1*^*-/-*^ and *Atg7*^*-/-*^;*Lkb1*^*-/-*^ mice sera compared with WT control mice. Data are mean ± s.e.m. *P<0.05, **P<0.01, ***P<0.001, and ****P<0.0001. b-d Metabolites that are significantly decreased in the sera of *Lkb1*^*-/-*^ and *Atg7*^*-/-*^;*Lkb1*^*-/-*^ mice compared with WT control mice at fasting state. Data are mean ± s.e.m. *P<0.05, ** P<0.01, ***P<0.001 and ****P<0.0001. e. Relative blood glucose levels of WT control, *Atg7*^*-/-*^, *Lkb1*^*-/-*^ and *Atg7*^*-/-*^;*Lkb1*^*-/-*^ adult mice normalized to WT control mice at fed state for the indicated time course after first TAM injection. Data are mean ± s.e.m. *P<0.05. f. Quantification of serum insulin levels of WT control, *Atg7*^*-/-*^, *Lkb1*^*-/-*^ and *Atg7*^*-/-*^;*Lkb1*^*-/-*^ adult mice at fasted state at 10 days post deletion. Data are mean ± s.e.m. *P<0.05, **P<0.01, ns: non-significant.

## Discussion

In this study, we demonstrated the intermingled essential and systemic roles of Lkb1 and autophagy in the maintenance of mouse homeostasis and survival via conditional whole-body deletion of *Lkb1* and *Atg7* in adult mice (Fig. 6). We found that acute Lkb1 loss led to damaged intestinal epithelium barrier and increased infection, and alteration in metabolic pathways necessary for maintaining host homeostasis, which was partially rescued by autophagy activation via inhibiting p53 induction. Thus, autophagy upregulation compensates for the acute Lkb1 loss to temporarily support the survival of adult mice.

**Fig 6.**
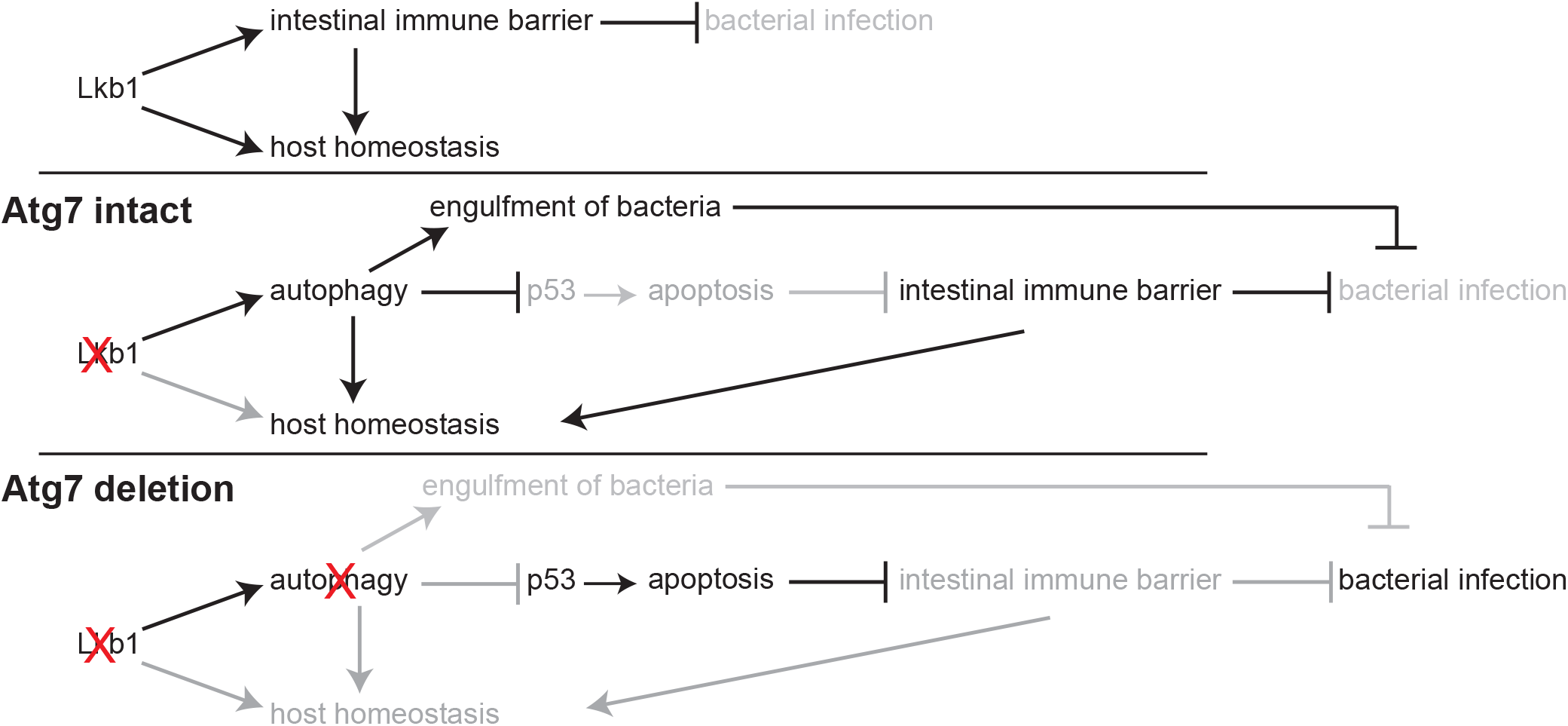
Mechanism by which Lkb1 interacts with autophagy to support adult mice homeostasis and survival. Loss of Lkb1 causes hypoglycemia, impaired intestinal epithelium barrier integrity, increased general infection, and disturbed host sera metabolism. When autophagy is intact, these dysfunctions are temporarily compensated by autophagy upregulation, partly through preventing p53 activation. However, autophagy deficiency further exacerbates the dysfunctions induced by *Lkb1* deletion, thereby accelerating mouse death.

An accumulating body of evidence suggests that Lkb1 phosphorylates AMPK and activates autophagy in response to energy crises (1-3, 18, 21). However, in addition to Lkb1, CaMKK, and TAK1 also trigger AMPK activation (22). Here we observed an upregulation of autophagy in *Lkb1*-deficient mice, suggesting that Lkb1-AMPK-mTORC1-autophagy axis may not be the only or the critical effector of Lkb1-mediated maintenance of adult mice homeostasis in our setting. This is consistent with a previous study that Lkb1-mediated energy metabolism is largely independent of Lkb1 regulation of AMPK and mTORC1 signaling in mouse hematopoietic stem cells (9). Moreover, AMPK and autophagy were activated in *Lkb1*-deficient lung tumors (19, 20).

Homozygous deletion of *Lkb1* in mouse is embryo-lethal (5), demonstrating an important role of Lkb1 in embryogenesis. Here, we reported an indispensable role of Lkb1 in maintaining the homeostasis and survival of adult mice. Mice carrying one inactivated allele of *Lkb1* (*Lkb1*^*+/-*^) recapitulate PJS and die at 11 months after birth due to the development of intestinal polyps (37). Hematopoietic stem cell-specific deletion of *Lkb1* led to limited mouse survival for up to 28 days due to pancytopenia (9). However, in our study, although mice died within 4 weeks after *Lkb1* ablation, no obvious reduction in different types of blood cells was observed. Instead, serum metabolomics analysis showed significant reduction in most of the essential and non-essential amino acids, certain metabolites related to the urea cycle and glycolysis in *Lkb1*^*-/-*^ mice compared with WT mice. Thus, the impaired systemic homeostasis by acute Lkb1 loss may be responsible for the death of *Lkb1*^*-/-*^ mice.

Lkb1 is involved in the development and maintenance of the goblet and Paneth cells (25). Specific deletion of *Lkb1* in intestinal stem cells leads to increased expression of pyruvate dehydrogenase kinase4 and reduced oxygen consumption, which reduces the population of stem cells and increase the levels of secretory cell number (24). The impaired goblet and Paneth cells are also characteristics of PJS in humans (37). Our whole-body deletion of *Lkb1* recapitulates the intestinal phenotype observed previously. Additionally, we found that the impaired intestinal structure was further exaggerated when *Atg7* is concurrently deleted with *Lkb1*. The major function of Goblet cells is attributed to the secretion of mucus to provide the epithelium immune barrier against bacterial invasion from the intestinal lumen (26). Loss of Lkb1 in intestinal-epithelium cells impaired intestinal barrier function (32). Here we demonstrated that the integrity of intestinal-epithelium barrier and function are damaged with a greater extent in *Atg7*^*-/-*^;*Lkb1*^*-/-*^ mice compared with *Lkb1*^*-/-*^ mice, leaving the mice vulnerable to infection. We found that broad-spectrum antibiotics supplementation could partially rescue the death of *Atg7*^*-/-*^;*Lkb1*^*-/-*^ mice, but not *Lkb1*^*-/-*^ mice. Although the role of autophagy in microbial defense is well established (38-40). Here we demonstrated for the first time the role of autophagy in preventing infection-related death induced by Lkb1 loss.

p53 is activated upon autophagy deletion, leading to increased apoptotic rates in mouse liver and brain, impairing the overall mouse homeostasis (36). A recent study shows that *Atg7* specific deletion in Lgr5^+^ epithelium cells promotes p53-induced apoptosis, leading to impaired integrity of intestinal barrier during stress (33). Our data was in line with these recent findings, where we found that whole body deletion of *p53* significantly rescued the survival of *Atg7*^*-/-*^;*Lkb1*^*-/-*^ mice. Conversely, *p53* deletion had no effect on the life span of *Lkb1*^*-/-*^ mice, showing that autophagy deletion in *Atg7*^*-/-*^;*Lkb1*^*-/-*^ mice affects mouse survival through activation of p53. Loss of intestinal barrier integrity is ascribed to the imbalance between cell proliferation in the crypt and cell migration towards the tip of villi where eventually cells undergo apoptosis (31). We observed increased apoptotic cell death in the villus tips of intestine in *Atg7*^*-/-*^;*Lkb1*^*-/*^ mice compared with *Lkb1*^*-/-*^ mice, which could be due to p53 activation in the absence of autophagy. However, the rate of cell proliferation in the crypt between *Atg7*^*-/-*^;*Lkb1*^*-/-*^ and *Lkb1*^*-/-*^ mice showed no significant difference. Thus, autophagy upregulation induced by acute Lkb1 loss plays an important role in maintaining a balance between cell proliferation and cell death in the intestine, presumably by inhibiting p53 induction.

Reduction of serum glucose and insulin levels were observed in both *Lkb1*^*-/-*^, and *Atg7*^*-/-*^;*Lkb1*^*-/-*^ mice, indicating that abnormal pancreatic function may lead to impaired gluconeogenesis and hypoglycemia. This data is also in parallel with muscle-specific *Lkb1*-deficient mice, which showed decreased blood glucose and insulin levels due to increased uptake of glucose through muscles (8). Most of the intermediates associated with amino acid metabolism, urea cycle, TCA cycle and glycolysis were significantly decreased by acute *Lkb1* ablation. Certain metabolites even showed higher extent of reduction in *Atg7*^*-/-*^;*Lkb1*^*-/-*^ mice compared with *Lkb1*^*-/-*^ mice. Moreover, the changes of metabolomics profiling in the intestine is consistent with that in serum, suggesting that this alteration occurs throughout the whole body, not a specific tissue. Taken together, both autophagy and Lkb1 are essential to maintain the host metabolism in adult mice.

## Material and Methods

### Mice

All animal experiments were performed in compliance with Rutgers Animal Care and Use Committee (IACUC) guidelines. Ubc-CreERT2 mice (17) (Jackson Laboratory) were cross-bred with *Atg7*^*flox/flox*^ mice (14), *Lkb1*^*flox/flox*^ mice (Jackson Laboratory) and p53^flox/flox^ mice (Jackson Laboratory) to generate *UbcCreERT2*^*/+*^;*Atg7*^*flox/flox*^ mice, *UbcCreERT2*^*/+*^;*Lkb1*^*flox/flox*^ mice, *UbcCreERT2*^*/+*^;*Atg7*^*flox/flox*^;*Lkb1*^*flox/flox*^ mice, *UbcCreERT2*^*/+*^;*Lkb1*^*flox/flox*^;*p53*^*flox/flox*^ mice and *UbcCreERT2*^*/+*^;*Atg7*^*flox/flox*^;*Lkb1*^*flox/flox*^;*p53*^*flox/flox*^ mice.

For the acute deletion of *Atg7, Lkb1* and *p53*, TAM (200 ul of suspended solution per 20 g body weight) was delivered to 8-10-week-old adult mice through intraperitoneal (IP) injections every 3 days for 4 times. In analysis of the survival curve, day 1 is the first day of TAM injection. Full deletion of the genes was obtained around day 9 (post-third injection).

To examine autophagy flux in adult mice, HCQ (100 mg per kg) was applied to the mice through IP injection.

To examine the effect of broad-spectrum antibiotics on the survival of *Lkb1*^*-/-*^ or *Atg7*^*-/-*^;*Lkb1*^*-/-*^ mice, broad-spectrum antibiotics Baytril (2.27% enrofloxacin) (5 mg per kg) was injected to the mice via IP twice per day.

### Serum assays

Blood glucose was measured using a True2Go glucose meter (Nipro Diagnostics), and liquid chromatography–mass spectrometry (LC-MS) metabolomics analysis (mentioned below). Serum insulin levels were assessed with an ultra-sensitive mouse insulin (Crystal Chem Inc., 90080) kit.

### Metabolomics analysis by LC-MS

Tissue or serum metabolites extracted using methanol:acetonitrile:water (40:40:20) (with 0.5% formic acid solution for tissue metabolite extraction and without formic acid for serum metabolite extraction) followed by neutralization with 15% ammonium bicarbonate were used for LC-MS, as described previously (13). Samples were subjected to reversed-phase ion-pairing chromatography coupled by negative mode electrospray ionization to a stand-alone orbitrap mass spectrometer (Thermo Fisher Scientific).

### Intestinal permeability analysis

In-vivo intestinal permeability was measured by FITC-dextran (Sigma Aldrich) gavage experiment. Mice were deprived from water overnight before oral gavaging with FITC-dextran at 44 mg per 100 g body weight. Subsequently, water was provided after gavage and blood samples were collected by cardiac puncture at 4 hours post-gavage. Sera were collected after centrifuging blood at 10,000 rpm for 10 minutes using 1.5 mL heparin-lithium coat tubes. Fluorescence intensity of the serum was measured, and the concentration of FITC-dextran was assessed according to the standard curve generated by the serial dilution of FITC-dextran (32).

### Histology and immunohistochemistry

Paraffin-embedded tissue sections were prepared as described previously (12) for H&E, and IHC staining. Antibodies utilized for IHC were Atg7 (Sigma Aldrich, A2856), Lkb1 (Santa Cruz Biotechnology, sc-32245), p62 (Enzo Life Sciences, PW9860-0100), LC3 (Nano Tools, LC3-5F10), Ki67 (Abcam, ab-15580), cleaved caspase-3 (Cell Signaling, 9661S), OLFM4 (Cell Signaling, 39141), lysozyme (Agilent, A0099), and p53 (Novus Biologicals, NB200-103SS). For the quantification of IHC on p53 and cleaved caspase-3, tissues were analyzed by quantifying at least 10 images at 20x magnification. A minimum of 200 cells were scored for each image.

### Western blotting

Tissues were snap-frozen in liquid nitrogen, ground using Cryomill in liquid nitrogen at 25Hz for 2 minutes, and then lysed in Tris lysis buffer (1M Tris-HCl, 1M NaCl, 0.1M EDTA, 10% NP40). Protein concentrations were measured using the Bio-Rad BCA reagent. Samples were probed with antibodies against Atg7 (Sigma Aldrich, A2856), Lkb1 (Santa Cruz Biotechnology, sc-32245), LC3 (Novus Biologicals, NB600-1384), p62 (American Research Products, 03-GP62-C), and β-actin (Sigma Aldrich, A1978). Western blots were quantified with Image J (National Institutes of Health). The intensities of bands were used to calculate relative ratios of the indicated protein over loading control (actin), which were then normalized based on the corresponding ratio in the wild type control sample.

### Real-time PCR

Total RNA was isolated from the tissues using Trizol (Invitrogen). cDNA was then reverse-transcribed from the total RNA using MultiScribe RT kit (Thermo Fisher). Real-time PCR was performed on Applied Biosystems StepOne Plus machine. *Cdkn1a (p21), PTEN* and *actin* genes were detected using predesigned commercial TaqMan primers for each gene accordingly (*Cdkn1a*: Mm00432448-m1, *PTEN*: mm00477210-m1, and *Actin*: Mm00607939-s1). Results were calculated using the ΔΔC_T_ method and then normalized to actin.

### Statistics

Data were expressed as the mean ± SEM. Statistical analyses were carried out with GraphPad Prism version 8.0 or Microsoft Excel. Significance in the Kaplan-Meier analyses to determine and compare the progression-free survival was calculated using the log-rank test. The mass spectra were analyzed by MAVEN software and the peak area of each detected metabolite was obtained. Statistical significance of metabolites was determined by a paired two-tailed Student’s t-test, and five mice from each genotype were used. P-value of <0.05 was considered statistically significant.

The heatmap of amino acids was generated by using R 3.6.1. program, and all values are processed by the mean normalization. Pearson algorithm was used for the hierarchical clustering of the rows.

## Abbreviations

LKB1: liver kinase B1
AMPK: 5’ adenosine monophosphate-activated protein kinase
mTORC1: mammalian target of rapamycin complex1
PJS: Peutz-Jeghers syndrome
ATG7: autophagy related7 gene
TCA: tricarboxylic acid
TAM: tamoxifen
UBC: ubiquitously expressed ubiquitin C
LC3: light chain3
TAK1: transforming growth factor beta-activated kinase1 (TAK1)
HCQ: hydroxychloroquine
WT: wild type
H&E: hematoxylin and eosin
IHC: immunohistochemistry
OLFM4: olfactomedin4
FITC: Fluorescein Isothiocyanate
CDKN1A: cyclin-dependent kinase inhibitor 1A
PTEN: phosphatase and tensin homolog

## Acknowledgments

We are grateful to Dr. Eileen White for her advice during the preparation of the manuscript; Wenping Wang in the Guo laboratory for helping with generation of heat map; Amy Lee, Nuha Syed, Akash Raju, Jerry Kong and Enrique Lopez in the Guo laboratory for helping with mouse ear tagging and genotyping. This work was supported by National Institute of Health (NIH) grant R01 CA237347-01A1, NIH grant K22 CA190521, American Cancer Society grant 134036-RSG-19-165-01-TBG, GO2 Foundation for Lung Cancer and the Lung Cancer Research Foundation to J.Y.G; New Jersey Commission on Cancer Research (NJCCR) grant DFHS18PPC021 to K.K; NJCCR grant DCHS19PPC013 to V.B; and NIH P30 CA072720 to Rutgers Cancer Institute of New Jersey.

## Author Contributions

J.Y.G. was the leading principal investigator who conceived and supervised the project. J.Y.G. and K.K. designed the experiments, interpreted the data and wrote the manuscript. K.K. performed the majority of the experiments and data analysis. V.B. performed the qRT-PCR and assisted with some of the survival experiments. Z.S.H. assisted with the mouse husbandry. S.F. assisted with tissue metabolite extractions. X.L. assisted with mouse genotyping.

## Supplementary figures

**Fig S1.**
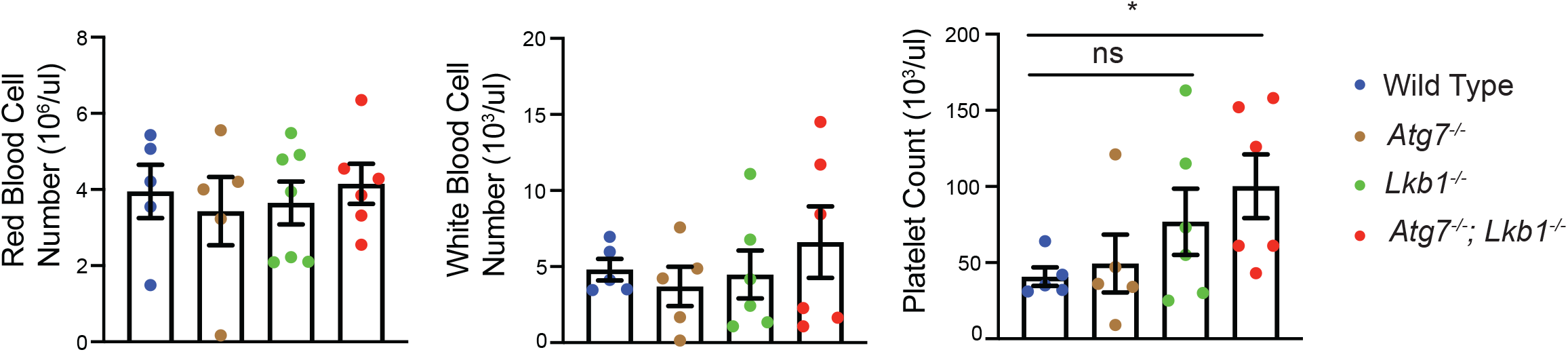
Autophagy and Lkb1 inhibitions do not lead to pancytopenia. Quantification of red blood cells, white blood cells, and platelets of WT control, *Atg7*^*-/-*^, *Lkb1*^*-/-*^, *and Atg7*^*-/-*^;*Lkb1*^*-/-*^ adult mice. Data are mean ± s.e.m. *P<0.05, ns: non-significant.

**Fig S2.**
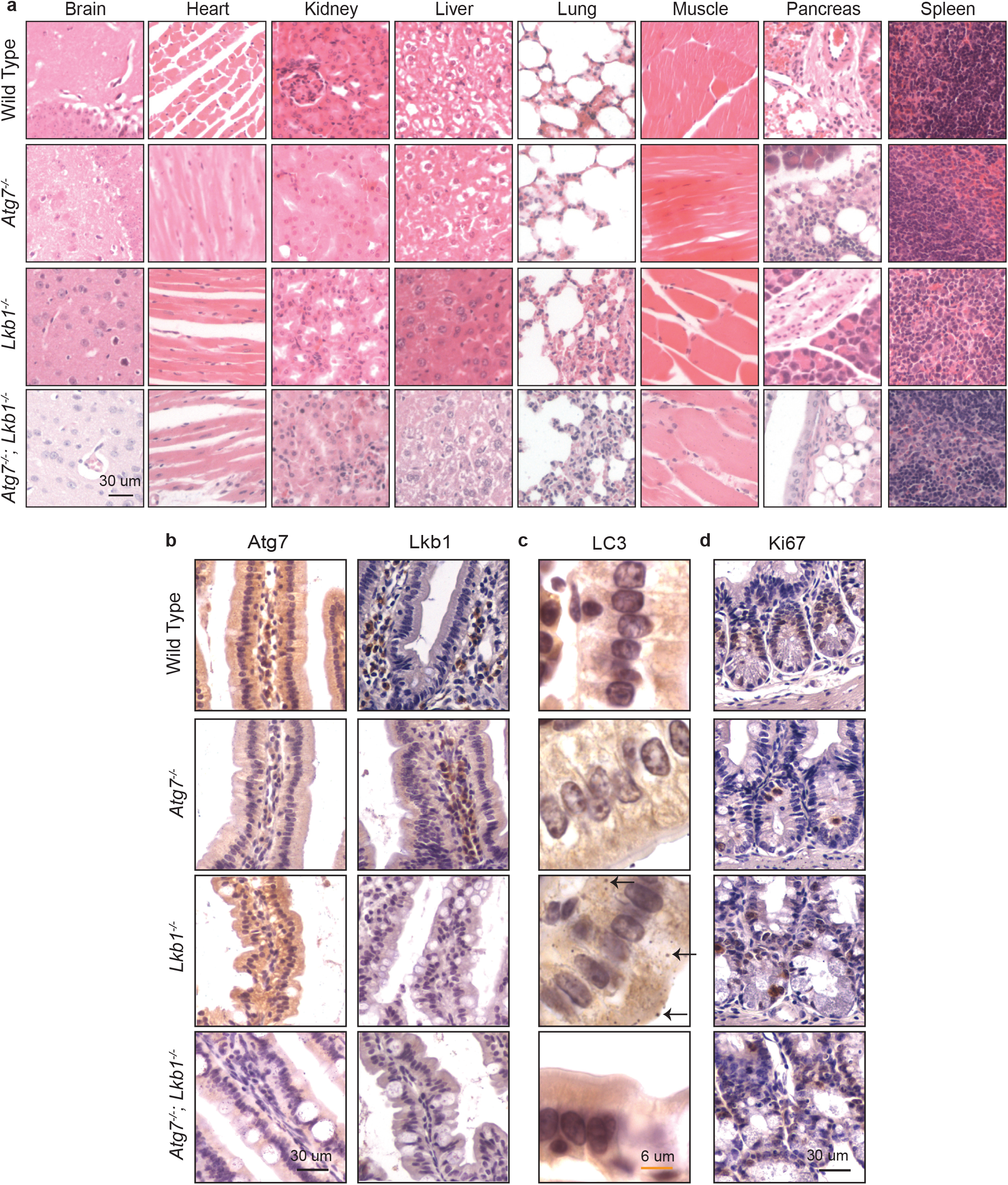
The histology of most mouse tissues and intestinal cell proliferation are not impacted by short-term deletion of Atg7 and Lkb1. a. Representative brain, heart, kidney, liver, lung, muscle, pancreas, and spleen histology (H&E staining) of WT control, *Atg7*^*-/-*^, *Lkb1*^*-/-*^, *and Atg7*^*-/-*^;*Lkb1*^*-/-*^ adult mice. b. Representative IHC for Atg7 and Lkb1 from intestine of WT control, *Atg7*^*-/-*^, *Lkb1*^*-/-*^ and *Atg7*^*-/-*^;*Lkb1*^*-/-*^ adult mice. c. Representative IHC for LC3 from intestine of WT control, *Atg7*^*-/-*^, *Lkb1*^*-/-*^ and *Atg7*^*-/-*^;*Lkb1*^*-/-*^ adult mice shows accumulation of LC3II puncta (arrows) in *Lkb1*-deficient mice. d. Representative IHC for Ki67 from intestine of WT control, *Atg7*^*-/-*^, *Lkb1*^*-/-*^ and *Atg7*^*-/-*^;*Lkb1*^*-/-*^ adult mice shows no significant difference in the rate of proliferation within the groups.

**Fig S3.**
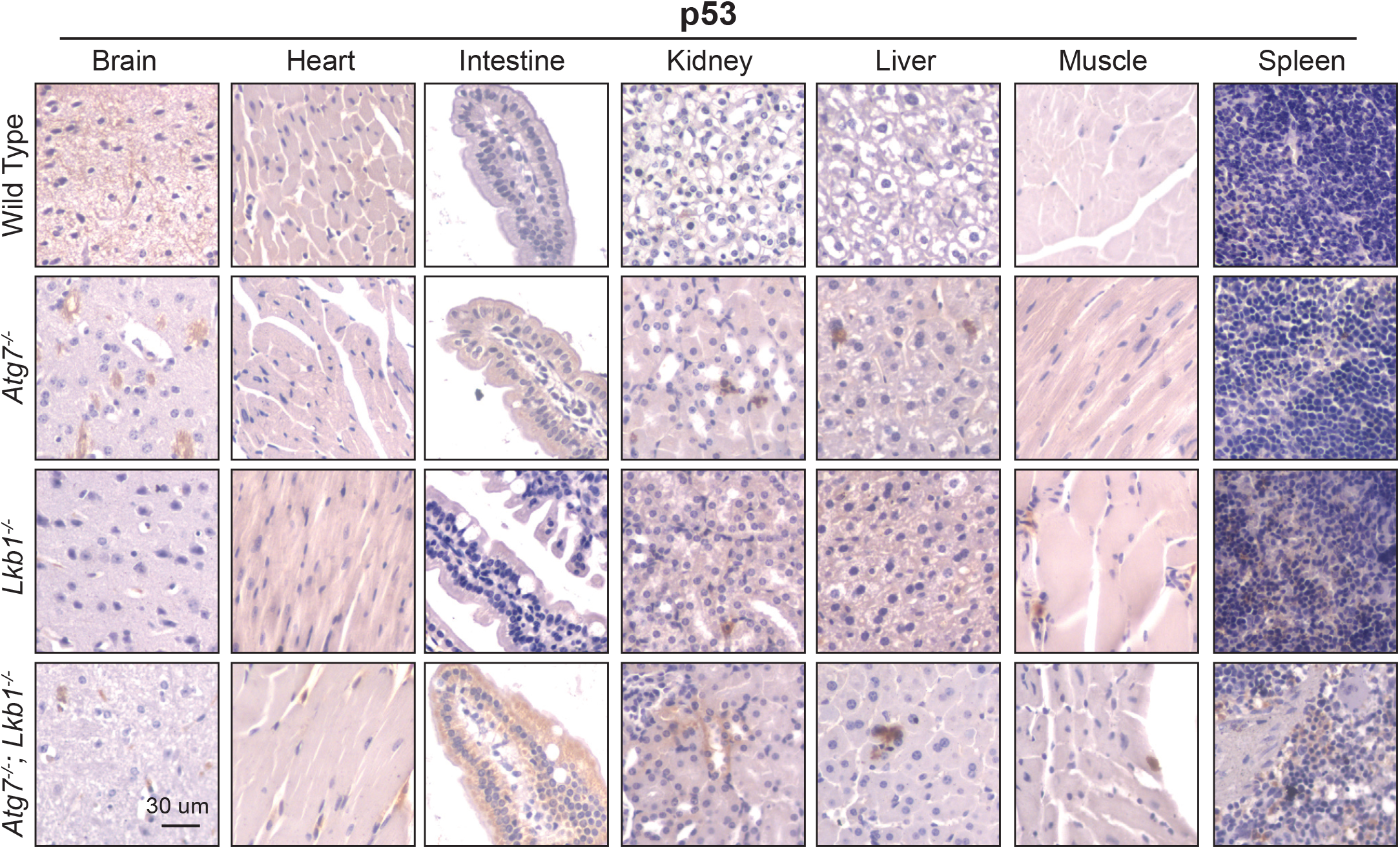
p53 is activated in the absence of autophagy. Representative IHC of p53 with lower magnification in brain, heart, intestine, kidney, liver, muscle and spleen of WT control, *Atg7*^*-/-*^, *Lkb1*^*-/-*^, *and Atg7*^*-/-*^;*Lkb1*^*-/-*^ adult mice shows the increase of nuclear p53 in *Atg7*-ablated tissues.

**Fig S4.**
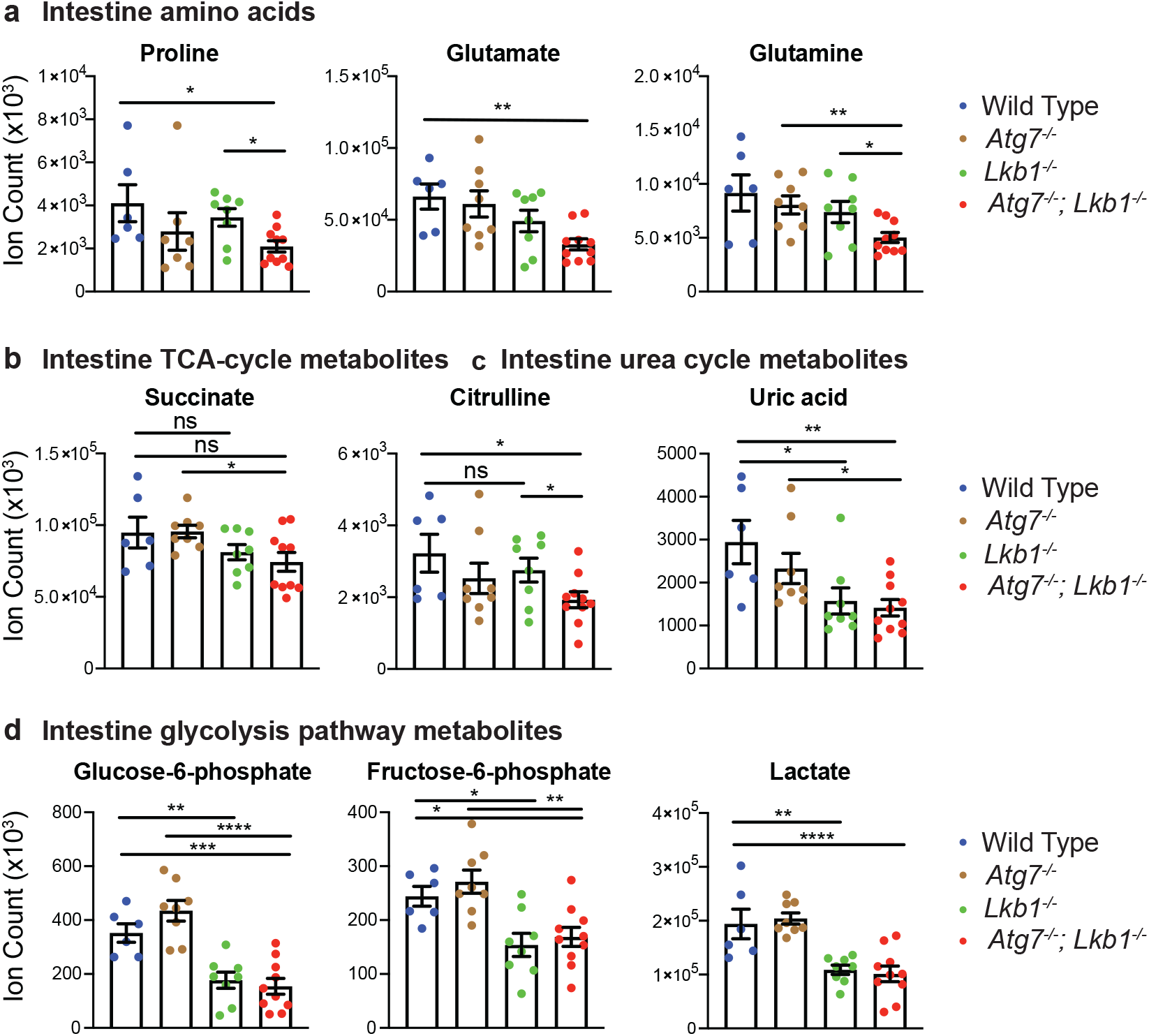
Acute loss of Lkb1 alone or together with Atg7 altered the levels of metabolites in intestine. a-d Metabolites that are significantly decreased in the intestine of *Lkb1*^*-/-*^ and *Atg7*^*-/-*^;*Lkb1*^*-/-*^ mice compared with WT control mice. Data are mean ± s.e.m. *P<0.05, **P<0.01, ***P<0.001, ****P<0.0001.

